# WITCH-NG: Efficient and Accurate Alignment of Datasets with Sequence Length Heterogeneity

**DOI:** 10.1101/2022.08.08.503232

**Authors:** Baqiao Liu, Tandy Warnow

**Affiliations:** Department of Computer Science, University of Illinois Urbana-Champaign, USA

## Abstract

Multiple sequence alignment (MSA) is a basic part of many bioinformatics pipelines, including in phylogeny estimation, prediction of structure for both RNAs and proteins, and metagenomic sequence analysis. Yet many sequence datasets exhibit substantial sequence length heterogeneity, both because of large insertions and deletions (indels) in the evolutionary history of the sequences and the inclusion of sequencing reads or incompletely assembled sequences in the input. A few methods have been developed that can be highly accurate in aligning datasets with sequence length heterogeneity, with UPP (Nguyen et al., 2015) one of the first methods to achieve good accuracy, and WITCH (Shen et al., Bioinformatics 2021) an improvement on UPP for accuracy, In this paper, we show how we can speed up WITCH. Our improvement includes replacing a critical step in WITCH (currently performed using a heuristic search) by a polynomial time exact algorithm using Smith-Waterman. Our new method, WITCH-NG (i.e., “next generation WITCH”, pronounced “witching”) achieves the same accuracy but is substantially faster. WITCH-NG is available in open source form at https://github.com/RuneBlaze/WITCH-NG.

## 1 Introduction

Multiple sequence alignment (MSA) is a fundamental task in computational biology and is a prerequisite for many downstream analyses such as phylogeny estimation [13], metagenomics [15, 20], and other applications. Over recent years, the assembly of large sequence datasets has led to the development of MSA methods that are able to scale to very large datasets (e.g., Clustal-Omega [23]) as well as techniques that use divide-and-conquer to maintain high accuracy (e.g., PASTA [11] and MAGUS [24]). Yet, accurate MSA estimation still remains challenging, especially under conditions such as high rates of evolution or sequence length heterogeneity.

Sequence length heterogeneity, in particular the presence of many short sequences, is a frequent characteristic of biological datasets (Figure 1 contains two examples). Sequence length heterogeneity can arise due to various reasons, such as the inclusion of many short reads or partially assembled sequences, or purely from evolutionary events such as domain-level deletions. MSA methods that are accurate on datasets without sequence length heterogeneity can degrade severely in accuracy under substantial presence of fragments, and the resulting subpar alignments will in-turn adversely affect downstream analyses [26]. Therefore, specialized MSA estimation methods that are robust to sequence length heterogeneity are valuable.

**Figure 1:**
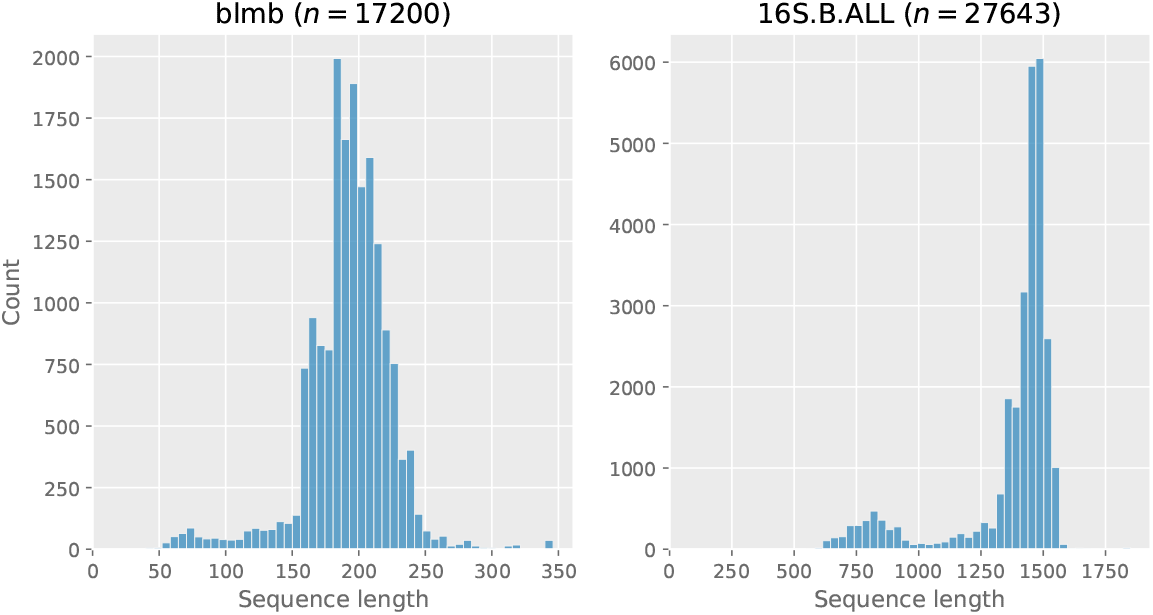
Sequence length histograms of two biological datasets that show sequence length heterogeneity. “blmb” comes from HomFam [23] and “16S.B.ALL” comes from the Comparative RNA Website (CRW) [3]. *n* denotes the number of sequences in the dataset.

One effective approach for aligning datasets with sequence length heterogeneity selects a set of “full-length” sequences from the input, aligns these sequences, and then adds the rest of the sequences (the “queries”) into the computed alignment. The last step where sequences are added into an existing alignment is its own bioinformatics problem. Notably, this last step of producing a extended alignment is a necessary step in other applications, including remote homology detection [16] and various tasks in metagenomics including taxon identification and abundance classification [10, 15, 20].

Methods that can produce this extended alignment include UPP [17], WITCH [21], MAFFT [7] (using the `--add` option), PaPaRa [1, 2], and HMMER [6] (using the popular `hmmbuild`+`hmmalign` pipeline).

WITCH is the most recent of the methods designed for this purpose, and has had the best accuracy of these methods. WITCH uses a two-stage technique to add sequences into a backbone alignment. In the first stage, it computes an ensemble of Hidden Markov Models (eHMM) to represent the backbone alignment, using tools from HMMER [4]. In the second stage, it adds the remaining sequences (i.e., the query sequences) into the backbone alignment, using the eHMM. At a. high level this two-stage structure is the same as in UPP, but WITCH executes the second stage differently. While UPP simply picks a single HMM from the eHMM to add a given query sequence into the backbone alignment, WITCH computes an extended alignment for the query sequence for each of the HMMs in the ensemble. It then combines these extended alignments, weighted by the probabilities it associates to each HMM in the ensemble, into a single extended alignment. This combination step is a “weighted consensus” of the alignments, and is performed using a graph algorithm called the “Graph Clustering Merger” (GCM) from MAGUS [24]. In other words, UPP uses a “mixture of experts” approach while WITCH uses an “ensemble” approach when combining information from the ensemble of HMMs.

The resulting pipeline improves the accuracy of UPP, an already accurate method, but at the expense of speed (Figure 2). Since WITCH and UPP differ only in the last step, this shows that the runtime cost is a result of its weighted consensus step where WITCH uses the top ten HMMs in the eHMM instead of just one, and more importantly because WITCH uses GCM, a general alignment merging method, for merging all the information in the different alignments (one for each HMM) in order to add the query sequence into the backbone alignment.

**Figure 2:**
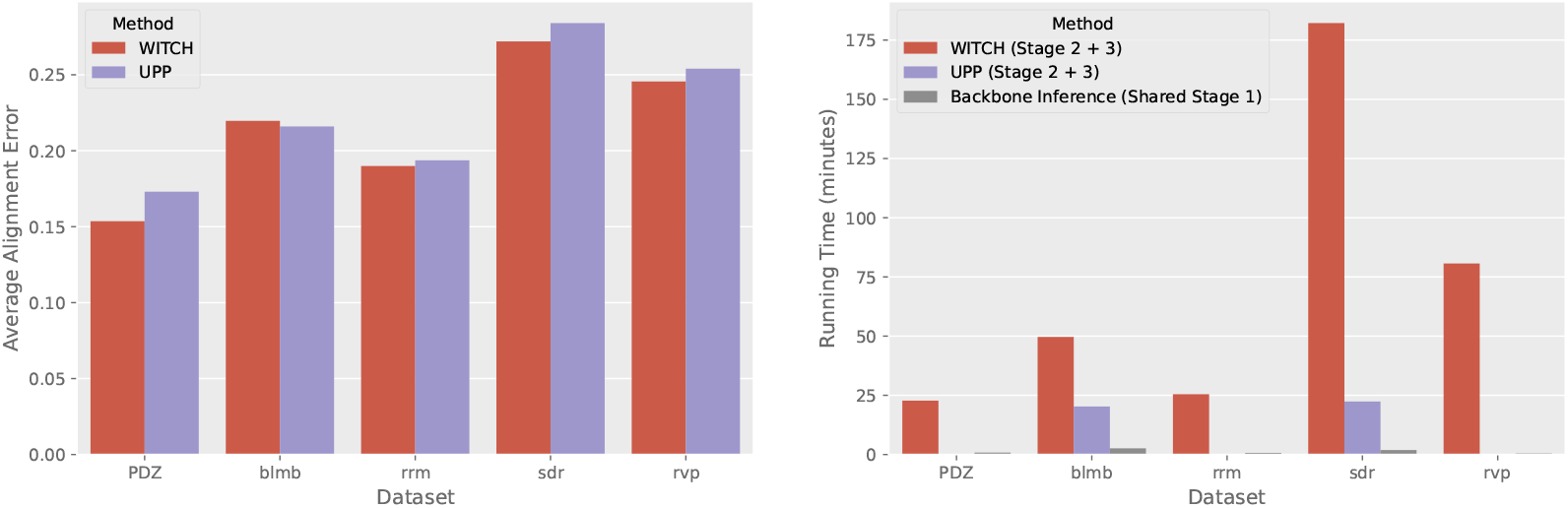
WITCH vs. UPP on five large Homfam datasets. Left: alignment error; right: runtime. As WITCH and UPP have the same stage 1 and nearly identical implementation for stage 2, this suggests that WITCH is substantially slower than UPP for stage 3. The datasets have respectively 14950, 17200, 27610, 50157, and 93681 sequences.

Here we present WITCH-NG, an algorithmic simplification of WITCH with an efficient implementation. We show that through simpler algorithmic design and better algorithmic engineering, WITCH-NG is a faster (and in many cases, much faster) version of WITCH, and in most cases nearly removes the running time penalty of using WITCH over UPP while retaining WITCH’s high accuracy. Therefore, WITCH-NG can be seen as an accurate (more accurate than UPP) alignment method with a slower yet comparable running time with UPP.

## 2 Background

### 2.1 WITCH

We present WITCH in detail, divided into three stages. The input is simply a set of unaligned sequences (with sequence length heterogeneity) and our goal is to produce an accurate alignment on such set of sequences. We also depict these stages in Figure 3.

**Figure 3:**
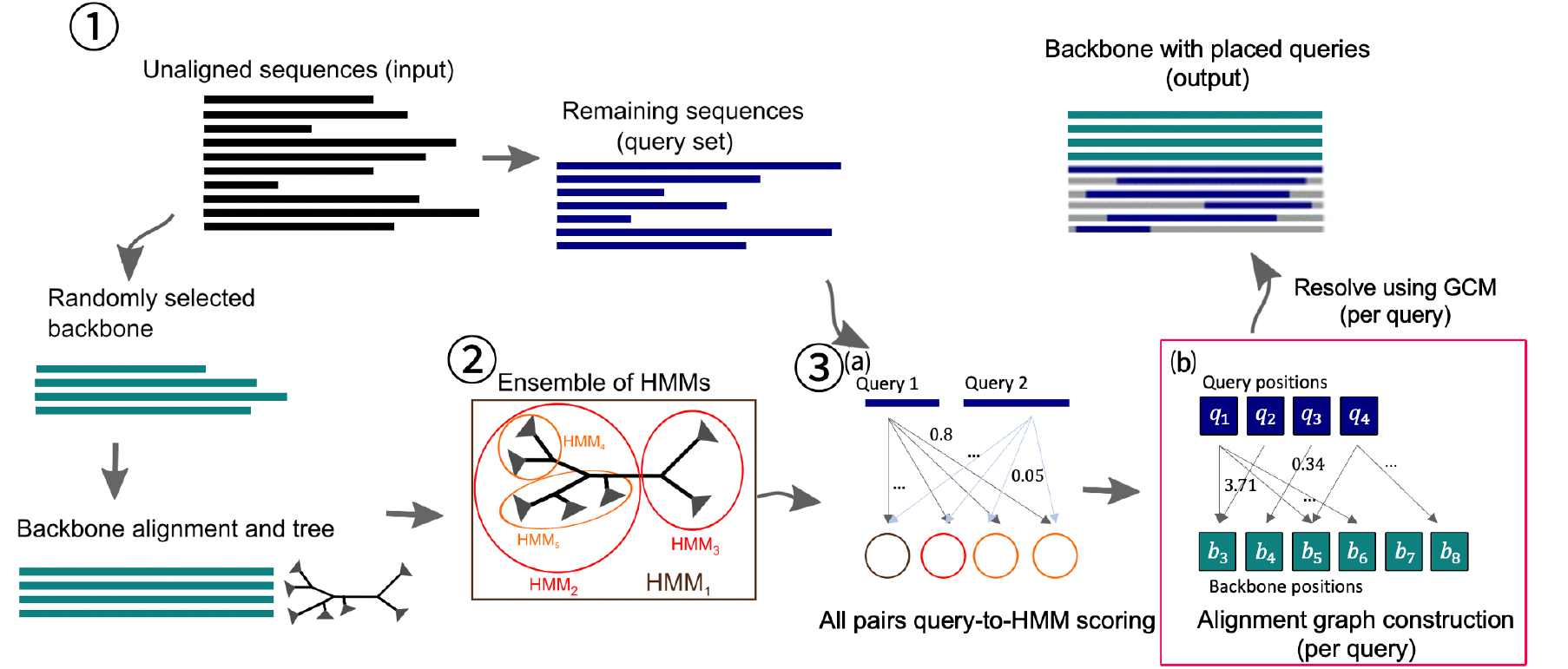
Overview of the original WITCH algorithm. Circled numbers show the stages of the algorithm, with “(a)” and “(b)” dividing Stage 3 into two steps. The red box on Stage 3(b) highlights that WITCH-NG in algorithm design only swaps out Stage 3(b). This figure is based on Figure 2 in [17], used under the Creative Commons Attribution (CC-BY) License.

#### 2.1.1 Stage 1: producing the backbone alignment and tree

The set of input sequences is first partitioned into two disjoint sets. The first set of “backbone sequences” is sampled from a full-length range, set as [0.75*l*, 1.25*l*] by default, where *l* is the median length of the input sequences. WITCH by default samples at most 1000 sequences from this range. The backbone sequences are then aligned by an existing accurate alignment method (current choice being MAGUS [24]), producing the “backbone alignment” *B*. The set of remaining sequences, which we call the “query set”, is denoted by *Q*. A tree *T* is estimated on *B*, with the current method of choice being the maximum likelihood heuristic FastTree2 [19].

#### 2.1.2 Stage 2: building the ensemble of HMMs

In this stage, *T* is used to hierarchically decompose *B* into overlapping subsets, upon which HMMs (using HMMER’s `hmmbuild` command) are built. Going into details, we define the size of a tree by its number of leaves and define the centroid edge of a tree by the edge that has the least difference in the number of leaves this edge separates. Then we recursively bisect *T* by the centroid edge until no more trees of size above *z* (*z* an algorithmic parameter, for our purposes *z* = 10) can be produced. All trees encountered throughout this recursion (including *T*) then defines a set of trees {*T*_*i*_}, which then in turn induces a set of subset alignments {*B*_*i*_}, where each *B*_*i*_ is *B* restricted to the taxon names of *T*_*i*_. A HMMER HMM is then built on each *B*_*i*_, resulting in the set of HMMs {*M*_*i*_}.

#### 2.1.3 Stage 3: placing the query sequences

This is the most involved stage. We divide this stage into two steps. In the first step (scoring), for each query *q* ∈ *Q* and for each model *M*_*i*_, a fitness score of *q* against *M*_*i*_, denoted as *w*_*q,i*_, is derived from the bitscore that HMMER’s `hmmsearch` produces. This fitness score (called “adjusted bitscore”) is derived under the assumption that each *q* is generated by exactly one of {*M*_*i*_} and *w*_*q,i*_ is the probability of *M*_*I*_ generating *q* (implying ∑_*i*_ *w*_*q,i*_ = 1 for any *q*). We now simply refer to *w*_*q,i*_ as the “weight” of *q* against *M*_*i*_.

In the second step (weighted consensus), each query *q* is independently placed into the backbone by constructing and resolving an “alignment graph”. Consider an weighted undirected bipartite graph *G* with nodes *q*1, *q*2, …, *q*_*m*_ where *m* is the length of the query sequence and nodes *b*1, *b*2, …, *b*_*n*_ where *n* is the number of columns in *B*. We add weighted edges to *G* as follows. First for computational efficiency, only the top *k* (by default *k* = 10) HMMs for each sequence by weight are retained. Let these HMMs be denoted as 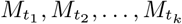. Intuitively, each of these HMM defines an optimal alignment (through `hmmalign`) of *q* against *B*_*i*_, hence also an optimal alignment of *q* against *B*. The homologies defined by all such HMMs can be represented as edges in *G* (each homology pair defines an undirected edge) while the edge weights are scaled by the weight of the HMMs. Concretely, if an 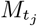 with weight *w* determines that character *x* of *q* and column *y* of *B* are homologous, an edge between *q*_*x*_ and *b*_*y*_ is added to *G* with edge weight *w c*_*y*_ where *c*_*y*_ is the number of non-gap characters in column *y* of *B*. GCM is then used as a blackbox weighted consensus method on *G* to resolve a merging between *q* and *B*. After each query sequence is placed into *B*, producing for each query sequence an “extended” alignment of *q* merged with *B*, the extended alignments are then merged transitively (i.e., if two query positions both map to the same backbone position, then these two query positions are also deemed homologous) into the final output alignment.

## 3 WITCH-NG: Redesign of WITCH

### 3.1 Algorithmic improvements

We modify the weighted consensus step (second step of the last stage, the step in the red box in Figure 3) of WITCH. Recall that at the second step of the last stage, we have already obtained weights *w*_*q,i*_ for each query sequence *q* and each HMM *M*_*i*_. In addition, for each query *q*, we have a construction for *G*, the alignment graph. Given that any weighted undirected bipartite graph on two components of size *m* and *n* can be naturally represented as a *m* × *n* matrix, we choose to represent *G* as this matrix denoted by *S* (i.e., *S*[*i, j*] is the weight between *q*_*i*_ and *b*_*j*_ in our above construction for *G*).

Critically, observe that *S* can be seen also as a scoring matrix for sequence alignment, where aligning position *i* of *q* against position *j* of *B* gets a reward of *S*[*i, j*]. WITCH-NG then sets all non-positive entries of *S* as -∞ (to forbid aligning two positions that are not supported by any of the HMMs) and runs a sequence alignment dynamic programming (DP) algorithm (say, Smith-Waterman [27]) to align *q* to *B* with a constant zero gap penalty (both Smith-Waterman and Needleman-Wunsch [14] simplify into the same DP algorithm under no gap penalty) using this *S* as the scoring matrix. The result is a placement of the query against the backbone. Given this way of placing queries, the rest of the algorithm identically follows that of WITCH. Each query is aligned to the backbone using this construction for the scoring matrix *S* and then merged transitively. Notice that the computationally intensive GCM blackbox is circumvented, replaced by a lightweight application of Smith-Waterman.

The problem of independently placing each query sequence into the backbone is thus a local alignment problem of aligning a query against the backbone, for which the classical approaches depend on scoring matrices and gap penalty schemes, and then use Smith-Waterman. The typical challenge in using this classical approach is to define the scoring scheme. However, in WITCH-NG, we use the matrix *S*[*i, j*] to define the scoring function, which makes this approach completely straightforward.

The reason this makes good sense is that Zaharias *et al*. [28] have argued that there is an implicit optimization problem in GCM (which is what WITCH does to compute the alignment of the query to the backbone), which is identical to the solution to the local alignment problem using this cost function. Specifically, GCM, although initial devised as a blackbox for alignment merging in MAGUS [25], has been shown to be a good heuristic for the optimization problem “Maximum Weight Trace for Alignment Merging (MWT-AM)” [29] (a straightforward generalization of the classical Maximum Weight Trace (MWT) problem [8] in bioinformatics). In short, MWT-AM defines an optimization criterion to the scenario when merging multiple disjoint “constraint” alignments (i.e., the homologies inside these alignments cannot be altered in the merging process) and when similarity scores between the constraint alignment columns have been obtained. Moreover, maximizing the MWT-AM score for merging many alignments is shown to be beneficial for alignment accuracy when using GCM [29].

Therefore, given how the original WITCH always uses GCM to merge two alignments, naturally we consider the problem of optimizing for the MWT-AM criterion (restricted to the case of two alignments). We give the definition of MWT-AM on two alignments below:

#### Definition 1

*(MWT-AM, trivial case) Given a weighted undirected bipartite graph with nodes q*1, …, *q*_*m*_ *and b*1, …, *b*_*n*_, *edges of form* (*q*_*i*_, *b*_*j*_) *with weight function w*((*q*_*i*_, *b*_*j*_)) *>* 0, *select a subset of edges T (the “trace”) maximizing*

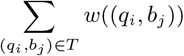

*subject to an additional “non-crossing” constraint, where for any two different* (*q*_*i*_, *b*_*j*_) *and* (*q*_*x*_, *b*_*y*_) *in T, either both i < x and j < y, or both x < i and y < j*.

A straightforward observation suggests that this trivial case is solvable by a simple DP algorithm (simplified version of either Smith-Waterman or Needleman-Wunsch without gap penalty), i.e., solvable in *O*(*mn*) time and *O*(*mn*) space, and this DP algorithm coincides with our presented algorithm on *S*. This observation is far from new and is mentioned in the paper introducing the Maximum Weight Trace (MWT) problem [8]. Therefore, *WITCH-NG can be seen as replacing a computationally intensive heuristic (i*.*e*., *GCM) to merge the query and the backbone by an exact polynomial time algorithm for the same problem*.

### 3.2 Implementation

We note two major differences in the implementation of WITCH-NG aiming for better running time compared to WITCH. Our first change is a strict improvement to the very last step of WITCH before the output, when the “extended alignments” (alignments only containing a query sequence and the backbone) are transitively merged. The original implementation in WITCH not only conceptually builds these extended alignments, but also writes these extended alignments to the disk verbatim before consuming them using a generic transitivity merging subroutine. This strategy is heavy in IO usage (writing extended alignments to disk then parsing them back into memory) and is extraneous because we only need to memorize each query letter’s matched position in the backbone in order to print the output alignment.

The second difference is that WITCH-NG implements a different strategy in invoking `hmmsearch` (the bottleneck step in obtaining the bitscores for UPP and in turn the weights for WITCH). WITCH-NG aggressively avoids invoking extra IO especially disk writes. Both WITCH and UPP use temporary files to divide up the query sequences into chunks for `hmmsearch` to consume and also to save outputs of `hmmsearch` across often times hundreds of HMMs. Combined with the IO usage of `hmmsearch` reading the HMMs (saved on disk), these steps might hinder efficient parallelization (i.e., operations can become IO-bound instead of CPU-bound). Ideally this problem is solved by moving on-disk structures into memory, e.g., saving the HMM structure in-memory if memory permits and then saving the query sequences in the native format used by HMMER in memory instead of having each `hmmsearch` instance repeating the reading and parsing of sequences from disk (then formatting its search results before us parsing the formatted results to extract the bitscores), but this solution requires serious implementation effort. As a makeshift solution, WITCH-NG relies on piping small chunks of sequences (saved in memory in plain text) to the `hmmsearch` command while also parsing the `hmmsearch` output in-memory in an attempt to minimize disk-write congestion. We do not claim this design as necessarily better than existing designs but more as a reasonable attempt to optimize such a common step (used in UPP, WITCH, HMMerge [18], etc., also in general used in any kind of HMMER-built database search against many sequences).

We finally note that although a sequence alignment DP algorithm is used as the engine for merging the query against the backbone. We in practice simply used a naive implementation (based on naive Smith-Waterman) despite many optimized variants of DP sequence alignment algorithms that are much faster on modern architectures (e.g., Farrar’s “striped Smith-Waterman” algorithm [5]).

## 4 Experimental Design

We assembled a diverse set of publicly available datasets (both simulated and biological, locations provided in the supplementary materials) to evaluate the difference of WITCH-NG and WITCH, along with WITCH’s predecessor UPP. The list of datasets and their respective statistics are shown in Table 4, where simulated conditions have been intentionally fragmented (1000M-HF series have half of the sequences fragmented to roughly 250bp in length [26]. RNASim-500bp has half of its sequences fragmented to an average length of 500bp [17], and when we later vary the backbone and query set size we directly assign full-length sequences to the backbone and the fragments to the query.)

We performed two experiments. In Experiment 1, we compare WITCH, WITCH-NG, and UPP on both simulated and biological nucleotide datasets (from the Comparative RNA Website (CRW) [3]) where we have reference alignments on the entire set of sequences. In Experiment 2, we compare WITCH, WITCH-NG, and UPP on the 10AA dataset (a selected set of ten protein alignment datasets that have curated alignments) [17] and also on the ten largest HomFam datasets [23] (up to 93681 sequences), which only have reference alignments on very small subsets of the input sequences.

### 4.1 Evaluation Criteria

Our goal is retain WITCH’s accuracy while improving speed, hence we track two sets of metrics. For accuracy, we use the Sum-of-Pairs (SP) error metrics computed by FastSP [12]. Briefly, each alignment encodes a set of homology pairs (defined by the columns of the alignment). SPFN (Sum-of-Pairs False Negative rate) is the proportion of true homology pairs (those from the reference alignment) that are not present in the estimated alignment, and 1 - SPFN equals the commonly used SP-score. SPFP (Sum-of-Pairs False Positive rate) is the proportion of estimated homology pairs not found in the reference, and 1 - SPFP is also known as the Modeler score. Both rates are defined from 0 to 1. For convenience, we often report the average of SPFN and SPFP, sometimes referred directly as the alignment error or average error. We evaluate the rates across the entire alignment (not just restricted on the query sequences) both to show the final accuracy of the methods as *de novo* alignment methods aligning sequences from scratch and to also take into account the homologies between query sequences and the backbone sequences (which will be ignored if only restricted to the query sequences).

For speed, we record the wall-clock running time of the methods assuming the backbone tree and backbone alignment are given (i.e., ignore the running time of Stage 1). Aside from the benefit of excluding a shared identical stage for all the methods, this wall-clock running time is exactly the running time for applications such as phylogenetic placement or metagenomics (e.g., taxon identification), where the reference alignment and tree often come prepackaged.

### 4.2 Computing Environment

All experiments were conducted on the Illinois Computing Cluster, a heterogeneous computing cluster where most runs are constrained to four hours with at least 64GB of available memory. For the most time-consuming dataset (16S.B.ALL), we explicitly did not constrain the running time to four hours. We ran all methods across 16 cores.

### 4.3 Other MSA Methods

We compare WITCH-NG to WITCH and UPP, each run in default mode, but all three methods used the same backbone alignment and eHMM. See Supplementary Materials for commands and version numbers. Methods other than UPP and WITCH were not selected for comparison as prior literature and preliminary results suggest that UPP is more accurate than other published methods (that we know of). For example, prior studies suggest UPP is more accurate than `hmmbuild+hmmalign` [17] and the MAFFT `--add` family of methods [22] (among those `--add` options that can scale to large datasets).

## 5 Results

### 5.1 Results for Experiment 1

Figure 4 presents an initial comparison of runtimes for WITCH and WITCH-NG on the RNASim-500bp datasets where we vary the size of the backbone and the number of query sequences. This comparison shows that WITCH is much faster than WITCH-NG across all settings, and that when there is a very large backbone (2000 sequences), WITCH can fail to complete due to runtime limitations when given many queries.

**Figure 4:**
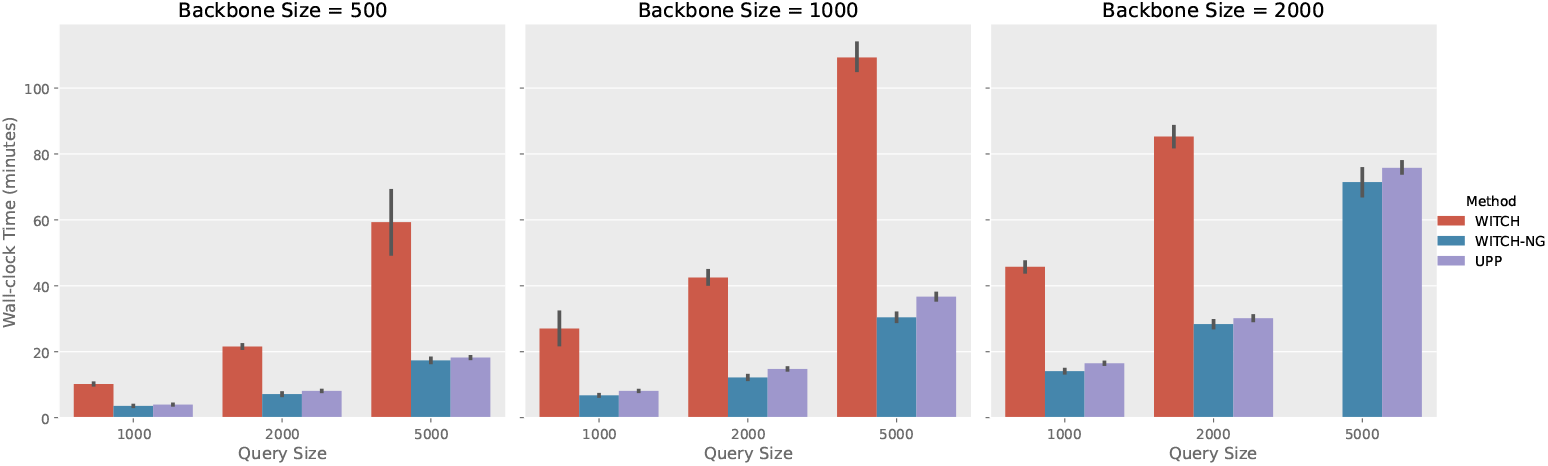
Wall-clock running time (minutes) between methods on the RNASim-500bp data, time measured for Stage 2 and Stage 3 (i.e., assuming backbone alignment and tree known). WITCH timed out on 1/5 replicates for backbone size 2000 and 5000 queries, and is excluded from the figure. All methods achieved roughly the same accuracy on these data (see Supplementary Material for the error rates).

While we just showed that WITCH-NG can be much faster than WITCH, WITCH-NG differs with WITCH in two sets of changes, algorithmic design and algorithmic engineering (implementation). It is important to confirm that the speed-up is not just due to better implementation. To isolate the impact of the two sets of changes, we created a version of WITCH’s code, where we replaced WITCH’s original GCM subroutine by a version of GCM that merges using our described variant of Smith-Waterman. This version of WITCH thus only includes the changes in algorithmic design. We then compare the speed of WITCH (the original implementation), “WITCH(Smith-Waterman)” (the modified WITCH just described), and WITCH-NG on two very different datasets used in this study, with the results shown in Figure 5. We see that the simplification in algorithm design is accountable for most (in the case of 1000M1-HF) and roughly half (for zf-CCHH) of the speed-up achieved by WITCH-NG, suggesting that both the design and engineering contributed to the speed-up, with perhaps the simplification in algorithm design contributing more.

**Figure 5:**
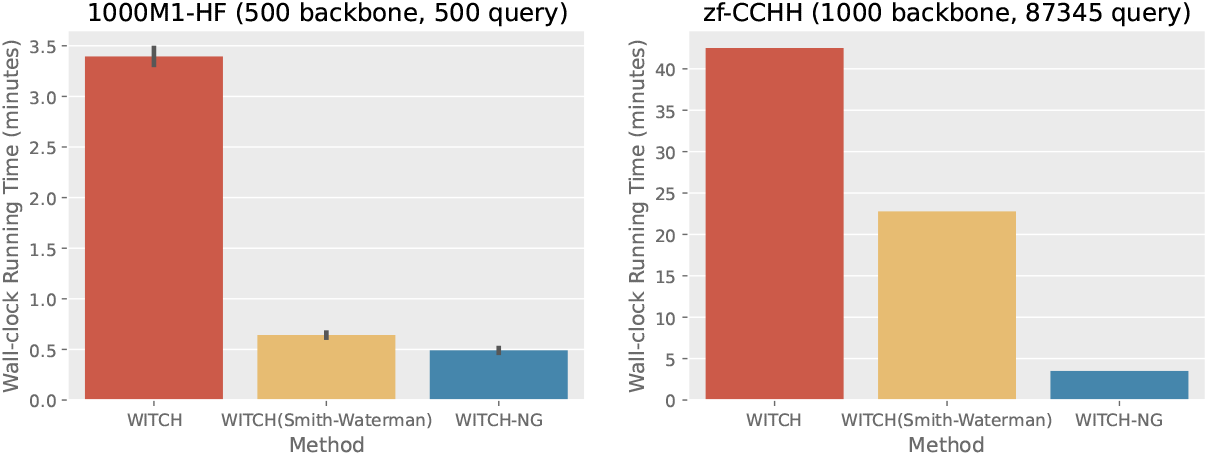
Exploring the effect of the algorithmic change (Smith-Waterman in lieu of default GCM) vs. the implementational changes, showing wall-clock running time (minutes, only measured on Stage 2 and 3) between methods on two very different data. “WITCH(Smith-Waterman)” denotes WITCH but with the GCM step replaced by the described Smith-Waterman step. On 1000M1-HF, the algorithmic change dominates the reduction in running time. On zf-CCHH, the implementational change is as important as the algorithmic change for the running time. On 1000M1-HF we show averages across replicates with standard error bars.

We show the results for Experiment 1 in both Table 2 (alignment error rates) and Figure 6 (left part, showing the running times). Looking at the average of SPFN and SPFP, WITCH and WITCH-NG are tied in being the most accurate method across all datasets, with the advantage over UPP most noticeable under higher levels of evolution (1000M2-HF and 1000M1-HF). Looking closer, WITCH and WITCH-NG have higher rates of false positive (SPFP) compared to UPP, but this disadvantage is more than compensated for by the lower SPFN of the two methods. WITCH-NG and WITCH have almost nonexistent differences in the error numbers, only on two cases differ by 0.001. As such, in later discussion, we sometimes refer to WITCH and WITCH-NG as one when there exists no apparent difference in their accuracy.

**Table 1:** Dataset statistics (average or range). ^1^ The 1000M series data have 20 replicates. 1000M1 has an outlier replicate as indicated in [24], hence removed. ^2^ RNASim-500bp has five replicates (on 5000 full-length sequences and 5000 fragmented sequences) and we sampled different configurations of backbone-query size to benchmark the algorithm. ^3^ HomFam datasets only have reference alignments on a small subset of the sequences. The last two of this row are derived from these small references. ^4^ Average p-dist (p-distance) is the proportion of homologous pairs of letters (as defined by the alignment) that are different, where a “homologous pair of letters” is two letters (nucleotides or amino acids) in the same column in the alignment.

| Dataset | # seqs. | Seq.<br>length | Align<br>length | p-dist.<sup>4</sup><br>(avg) |
| --- | --- | --- | --- | --- |
| Simulated Nucleotide |  |  |  |  |
| 1000M1-HF <sup>1</sup> | 1000 | 631.3 | 3960 | 0.694 |
| 1000M2-HF | 1000 | 634.3 | 3972 | 0.683 |
| 1000M3-HF | 1000 | 629.6 | 2723 | 0.660 |
| 1000M4-HF | 1000 | 629.6 | 2571 | 0.495 |
| RNASim-500bp <sup>2</sup> | 1500-7000 | 1025.4 | 21946 | 0.408 |
| Nucleotide |  |  |  |  |
| 5S.3 | 5507 | 105.6 | 414 | 0.418 |
| 5S.T | 5751 | 106.2 | 436 | 0.425 |
| 16S.3 | 6323 | 1557.2 | 8716 | 0.315 |
| 16S.T | 7350 | 1492.1 | 11856 | 0.345 |
| 16S.B.ALL | 27643 | 1371.9 | 6857 | 0.210 |
| Protein |  |  |  |  |
| 10AA | 303-807 | 432.7 | 1745.3 | 0.671 |
| Homfam<sup>2</sup> | 14950-93681 | 149.8 | 273.5 | 0.690 |

**Table 2:** Alignment error rates on nucleotide datasets (four simulated datasets and four biological datasets) placing into the same backbone alignment and tree. SPFN and SPFP are alignment error rates (lower is better) defined in text, and “Avg. Error” is the average of these two values. Best values (with ties within 0.001) are boldfaced.

| Dataset | Method | SPFN | SPFP | Avg. Error |
| --- | --- | --- | --- | --- |
| 1000M4-HF | WITCH | **0.014** | **0.010** | **0.012** |
|  | WITCH-NG | **0.014** | **0.010** | **0.012** |
|  | UPP | 0.016 | **0.010** | **0.013** |
| 1000M3-HF | WITCH | **0.048** | **0.039** | **0.043** |
|  | WITCH-NG | **0.048** | **0.039** | **0.043** |
|  | UPP | 0.054 | **0.038** | 0.046 |
| 1000M2-HF | WITCH | **0.128** | **0.106** | **0.117** |
|  | WITCH-NG | **0.128** | **0.106** | **0.117** |
|  | UPP | 0.137 | **0.106** | 0.121 |
| 1000M1-HF | WITCH | **0.172** | **0.143** | **0.157** |
|  | WITCH-NG | **0.171** | **0.143** | **0.157** |
|  | UPP | 0.182 | **0.142** | 0.162 |
| 5S.3 | WITCH | **0.089** | **0.086** | **0.088** |
|  | WITCH-NG | **0.089** | **0.086** | **0.088** |
|  | UPP | 0.093 | **0.085** | **0.089** |
| 5S.T | WITCH | **0.117** | **0.104** | **0.110** |
|  | WITCH-NG | **0.117** | **0.104** | **0.111** |
|  | UPP | 0.120 | **0.103** | 0.112 |
| 16S.3 | WITCH | **0.089** | **0.166** | **0.128** |
|  | WITCH-NG | **0.089** | **0.166** | **0.128** |
|  | UPP | **0.090** | **0.166** | **0.128** |
| 16S.T | WITCH | **0.172** | 0.177 | **0.175** |
|  | WITCH-NG | **0.172** | 0.178 | **0.175** |
|  | UPP | 0.189 | **0.171** | 0.180 |
| 16S.B.ALL | WITCH | **0.044** | **0.045** | **0.044** |
|  | WITCH-NG | **0.044** | **0.045** | **0.044** |
|  | UPP | **0.045** | **0.045** | **0.045** |

**Figure 6:**
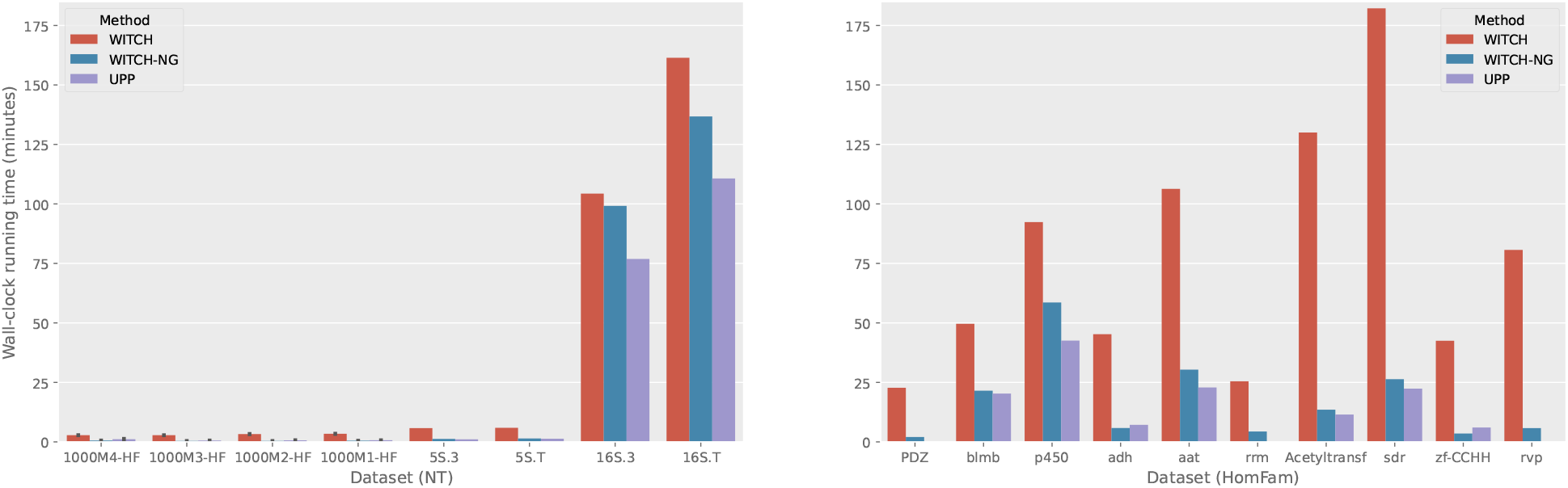
Wall-clock running time (minutes) for Experiments 1 and 2, time measured for Stage 2 and Stage 3 (i.e., assuming backbone alignment and tree known), 16S.B.ALL is excluded from the figure due to high running time of all methods (WITCH took 11 h 17 min, WITCH-NG took 8 h 4 min, and UPP took 5 h 18 min). The included NT datasets have 1000 sequences for the “HF” conditions shown, of which 500 are in the backbone and 500 are in the query. The rest (from left to right) have 5507, 5751, 6323, and 7350 sequences respectively. The HomFam datasets (right subfigure) have respectively 14950, 17200, 21013, 21331, 25100, 27610, 46285, 50157, 88345, and 93681 sequences. All data except the “HF” conditions have 1000 backbone sequences and the rest are included in the query.

For running time, we see different trends for the 16S series of the CRW data and the rest of the nucleotide datasets. For the four simulated conditions (1000M-HF) and the 5S series, WITCH-NG is much faster than WITCH (by multiple factors), and close to the runtime of UPP (typically a bit slower, but occasionally faster). On all the 16S datasets, WITCH-NG is faster than WITCH, but by a smaller amount than on the other datasets, and is slower than UPP. Thus, WITCH-NG speeds up WITCH on all datasets, sometimes by a great margin, and (except for the 16S datasets) is reasonably close to UPP in runtime.

### 5.2 Results for Experiment 2

On protein datasets, for 10 AA, all methods have near identical averaged alignment error as seen in Table 3. It still can be seen that WITCH(-NG) trades off SPFP for SPFN compared to UPP, but in this case not resulting in lower average error. All methods finished under one minute for these datasets with no notable difference in running time; hence we omit showing the running time results.

**Table 3:** Alignment error rates on 10AA and Homfam (averaged across the ten datasets in each collection). SPFN and SPFP are alignment error rates (lower is better) defined in text, and “Avg. Error” is the average of these two values. Best values (with ties within 0.001) are boldfaced.

| Dataset | Method | SPFN | SPFP | Avg. Error |
| --- | --- | --- | --- | --- |
| 10AA | WITCH | **0.233** | 0.189 | **0.211** |
|  | WITCH-NG | **0.233** | 0.189 | **0.211** |
|  | UPP | 0.236 | **0.185** | **0.210** |
| Homfam | WITCH | **0.327** | 0.124 | **0.225** |
|  | WITCH-NG | **0.327** | 0.125 | **0.226** |
|  | UPP | 0.336 | **0.122** | 0.229 |

On the ten largest HomFam datasets (second row, Table 3), WITCH and WITCH-NG have practically no difference in accuracy but are both more accurate than UPP. Across individual datasets in HomFam (shown in Supplementary Material), we see more mixed signals in terms of accuracy, with WITCH(-NG) in general better in average error than UPP (in five out of ten datasets). On two datasets UPP is better than WITCH(-NG) (on blmb and Acetyltransf) and in other cases tied. Interestingly, we see cases where WITCH and WITCH-NG differ more in the error metrics, for example on Acetyltransf where WITCH-NG is 0.01 point better in SPFP and on sdr where WITCH is almost 0.01 point better in SPFP. The higher variance in accuracy (compared to the previous experiment) might have come from the fact that HomFam datasets have very few reference sequences (for example, Acetyltransf has reference alignment on 6 sequences out of 46285 unaligned sequences).

For running times on HomFam (right subfigure, Figure 6), we see a dramatic reduction of the running time from WITCH to WITCH-NG in many cases, especially in cases where the runtime penalty of WITCH compared to UPP is the most obvious. Noticeably, WITCH-NG has running time quite close to UPP across all HomFam datasets, in most cases only paying a small penalty in running time compared to UPP and in two cases even faster than UPP. We conclude that for this type of large datasets (1000 backbone and more than 10000 query sequences), WITCH-NG dramatically speeds up WITCH with little difference in accuracy, achieves running time close to UPP and is more accurate than UPP.

## 6 Discussion

The speed-up of WITCH-NG over WITCH varies across datasets (in particular, the 16S series are outliers). As seen in Figure 4, the number of sequences in the input cannot explain this difference well, as WITCH-NG is dramatically faster than WITCH on the dataset with the largest number of sequences, but is only slightly faster on some smaller datasets. One potential explanation is sequence length, as the 16S series have the longest sequence lengths, and many long sequences lead to HMMs with many states, which increases the time for Stage 3(a). When Stage 3(a) uses much of the runtime, WITCH-NG will not differ much from WITCH in runtime, as the methods share largely the same algorithm for Stage 3(a). However, future work is needed to understand this variability.

## 7 Conclusion

WITCH-NG is an accurate multiple sequence alignment method designed for datasets with sequence length heterogeneity. WITCH-NG improves on the speed of WITCH, as a result of its algorithmic simplification and better implementation, the current most accurate method for this problem, and has running time comparable to that of UPP.

Although WITCH-NG is designed for *de novo* multiple sequence alignment, it can also be used directly to add sequences into alignments, a problem that arises in updating existing alignments and trees as new sequences are assembled (e.g., in phylogenetic placement [9]) and in microbiome analysis [15]. WITCH-NG should be evaluated for use in these applications, especially for those analyses where runtime is extremely important (e.g., taxon identification in metagenomics). Additional improvements to runtime and memory usage are also possible and should be investigated.

## 8 Acknowledgments

This work is supported in part by funds from the National Science Foundation (NSF: 2006069). It was also supported by the U.S. Department of Energy, Office of Science, Office of Biological and Environmental Research under the Secure Biosystems Design Initiative and by the Laboratory Directed Research and Development (LDRD) program of Sandia National Laboratories, which is a multimission laboratory managed and operated by National Technology and Engineering Solutions of Sandia, LLC, a wholly owned subsidiary of Honeywell International Inc, for the U.S. Department of Energy’s National Nuclear Security Administration under contract DE-NA0003525.

