## Supplementary Material for "WITCH-NG: Efficient and Accurate Alignment of Datasets with Sequence Length Heterogeneity"

August 8, 2022

**3 Dataset Availability** **6**

### 1 Additional Tables

| Backbone Size | Query Size | Method | SPFN | SPFP | Avg. Error |
|---|---|---|---|---|---|
| 500 | 1000 | WITCH | **0.100** | **0.097** | **0.099** |
| | | WITCH-NG | **0.100** | **0.097** | **0.099** |
| | | UPP | 0.102 | **0.097** | **0.099** |
| | 2000 | WITCH | **0.099** | **0.094** | **0.097** |
| | | WITCH-NG | **0.099** | **0.095** | **0.097** |
| | | UPP | 0.102 | **0.094** | **0.098** |
| | 5000 | WITCH | **0.099** | **0.093** | **0.096** |
| | | WITCH-NG | **0.098** | **0.093** | **0.096** |
| | | UPP | 0.102 | **0.093** | 0.097 |
| 1000 | 1000 | WITCH | **0.096** | **0.093** | **0.095** |
| | | WITCH-NG | **0.096** | **0.093** | **0.095** |
| | | UPP | **0.097** | **0.093** | **0.095** |
| | 2000 | WITCH | **0.096** | **0.091** | **0.094** |
| | | WITCH-NG | **0.096** | **0.092** | **0.094** |
| | | UPP | 0.098 | 0.091 | **0.094** |
| | 5000 | WITCH | **0.096** | **0.090** | **0.093** |
| | | WITCH-NG | **0.095** | **0.090** | **0.093** |
| | | UPP | 0.098 | **0.090** | **0.094** |
| 2000 | 1000 | WITCH | **0.093** | **0.088** | **0.091** |
| | | WITCH-NG | **0.093** | **0.088** | **0.091** |
| | | UPP | **0.094** | **0.088** | **0.091** |
| | 2000 | WITCH | **0.093** | **0.088** | **0.090** |
| | | WITCH-NG | **0.093** | **0.088** | **0.090** |
| | | UPP | **0.094** | **0.087** | **0.091** |
| | 5000 | WITCH-NG | **0.092** | **0.087** | **0.089** |
| | | WITCH | X | X | X |
| | | UPP | 0.094 | **0.086** | **0.090** |

Table 1: Alignment error rates on RNASim-500bp, averaged over five replicates. WITCH timed out on 1/5 replicates on the 200 backbone 5000 query configuration, hence excluded and marked with "X". SPFN (sum-of-pairs false negative) refers to the proportion of pairwise homologies in the reference alignment missing in the estimated alignment, SPFP (sum-of-pairs false positive) refers to the proportion of homologies in the estimated alignment that are not in the reference alignment, and "Avg. Error" is the average of these two values. Best values (with ties within 0.001) are boldfaced.

| Dataset | Method | SPFN | SPFP | Avg. Error |
|---|---|---|---|---|
| PDZ | WITCH | **0.211** | **0.096** | **0.154** |
| | WITCH-NG | **0.211** | **0.096** | **0.154** |
| | UPP | 0.239 | 0.107 | 0.173 |
| blmb | WITCH | **0.300** | 0.139 | 0.220 |
| | WITCH-NG | **0.300** | 0.142 | 0.221 |
| | UPP | 0.310 | **0.122** | **0.216** |
| p450 | WITCH | 0.279 | 0.183 | **0.231** |
| | WITCH-NG | **0.277** | 0.186 | **0.231** |
| | UPP | 0.298 | **0.175** | 0.237 |
| adh | WITCH | **0.638** | **0.003** | **0.321** |
| | WITCH-NG | **0.638** | **0.003** | **0.321** |
| | UPP | 0.641 | **0.003** | **0.322** |
| aat | WITCH | **0.182** | **0.106** | **0.144** |
| | WITCH-NG | **0.182** | **0.106** | **0.144** |
| | UPP | **0.183** | **0.106** | **0.145** |
| rrm | WITCH | **0.246** | **0.134** | **0.190** |
| | WITCH-NG | **0.247** | 0.136 | **0.191** |
| | UPP | 0.253 | **0.134** | 0.194 |
| Acetyltransf | WITCH | **0.548** | 0.167 | 0.358 |
| | WITCH-NG | **0.548** | 0.157 | 0.352 |
| | UPP | 0.550 | **0.146** | **0.348** |
| sdr | WITCH | **0.404** | **0.140** | **0.272** |
| | WITCH-NG | **0.405** | 0.148 | 0.276 |
| | UPP | 0.421 | 0.147 | 0.284 |
| zf-CCHH | WITCH | **0.186** | **0.055** | **0.120** |
| | WITCH-NG | **0.186** | **0.055** | **0.120** |
| | UPP | **0.186** | **0.056** | **0.121** |
| rvp | WITCH | **0.273** | **0.218** | **0.246** |
| | WITCH-NG | **0.273** | **0.218** | **0.246** |
| | UPP | 0.281 | 0.227 | 0.254 |
| Average across ten | WITCH | **0.327** | 0.124 | **0.225** |
| | WITCH-NG | **0.327** | 0.125 | **0.226** |
| | UPP | 0.336 | **0.122** | 0.229 |

Table 2: Alignment error rates on the ten largest HomFam datasets. SPFN (sum-of-pairs false negative) refers to the proportion of pairwise homologies in the reference alignment missing in the estimated alignment, SPFP (sum-of-pairs false positive) refers to the proportion of homologies in the estimated alignment that are not in the reference alignment, and "Avg. Error" is the average of these two values. Best values (with ties within 0.001) are boldfaced.

| **Name** | **Type** | **# seqs.** | **Seq. length** | **Align length** | **% gappy** | **p-dist.** | |
|---|---|---|---|---|---|---|---|
| | | | | | | **avg.** | **max** |
| Simulated | | | | | | | |
| 1000M1-HF | Sim NT | 1000 | 631.3 | 3960 | 84.0 | 0.694 | 1.0 |
| 1000M2-HF | Sim NT | 1000 | 634.3 | 3972 | 83.8 | 0.683 | 1.0 |
| 1000M3-HF | Sim NT | 1000 | 629.6 | 2723 | 76.7 | 0.660 | 1.0 |
| 1000M4-HF | Sim NT | 1000 | 629.6 | 2571 | 75.3 | 0.495 | 1.0 |
| RNASim-LF | Sim NT | 1500-7000 | 1025.44 | 21946 | 95.3 | 0.408 | 1.0 |
| Nucleotide | | | | | | | |
| 5S.3 | Bio NT | 5507 | 105.6 | 414 | 74.5 | 0.418 | 1.0 |
| 5S.T | Bio NT | 5751 | 106.2 | 436 | 75.6 | 0.425 | 1.0 |
| 16S.3 | Bio NT | 6323 | 1557.2 | 8716 | 82.1 | 0.315 | 0.833 |
| 16S.T | Bio NT | 7350 | 1492.1 | 11856 | 87.4 | 0.345 | 0.901 |
| 16S.B.ALL | Bio NT | 27643 | 1371.9 | 6857 | 80.0 | 0.210 | 0.769 |
| Protein | | | | | | | |
| 10AA | Bio AA | 303-807 | 432.7 | 1745.3 | 60.0 | 0.671 | 0.936 |
| Ten largest Homfam | Bio AA | 14950-93681 | 149.8 | 273.5 | 22.8 | 0.690 | 0.823 |

Table 3: Dataset statistics expanded. For HomFam, the last four columns are derived from the small reference alignments.

### 2 Software Versions and Commands

#### 2.1 Methods for Adding Queries

These methods assume the existence of the following three files:

- `$bb_aln`, (path to) the backbone alignment
- `$bb_tre`, the backbone tree
- `$q`, the query sequences

In addition, WITCH and UPP requires specifying the "molecule" type of the input, either "`dna`", "`rna`", or "`amino`". Let `$molecule` be this variable. Both WITCH and UPP requires a working directory, which we denote by `$work_dir`. The number 16 that occurs across the commands is the number of threads we used.

We ran UPP version 4.5.2, with the following command:

```
python3 run_upp.py -m $molecule -x 16 -s $q -a $bb_aln -t $bb_tre -d $work_dir -o
    $output_suffix
```

We ran WITCH version 0.2.1, with the following command:

```
python3 witch.py --molecule $molecule -t 16 -q $q -b $bb_aln -e $bb_tre -d $work_dir
```

We ran WITCH-NG version 0.0.2, with the following command:

```
witch-ng add --threads 16 -i $q -b $bb_aln -t $bb_tre -o $output_path
```

#### 2.2 HMMER Commands

We invoke HMMER (version 3.1b2 to keep it consistent with WITCH) inside WITCH-NG as subprocesses. The commands are given below:

```
hmmbuild --cpu 0 --informat afa --ere 0.59 --symfrac 0.0 $hmm_outpath -
hmmsearch --cpu 0 --noali --max -E 999999999 $hmm_path -
hmmalign --informat fasta --outformat afa $hmm_path -
```

#### 2.3 Other Commands

For computing the alignment error statistics, we used FastSP [2] (v1.7.1) with the following command:

```
java -jar FastSP.jar -ml -e $est -r $ref
```

For the "WITCH(Smith-Waterman)" method, we used a custom version of WITCH, located here (note that this link does not point to the official version of WITCH, but instead points to a branch with the original GCM swapped out), with the exact same commands used for WITCH listed above.

For aligning the backbone alignment and to infer trees on the backbone alignment, we either directly ran WITCH (which will as a first step split the dataset and align the backbone alignment, then inferring a tree on the backbone alignment) terminating it once the backbone alignment and tree is produced, or we manually align the backbone sequences and infer trees on the backbone alignment using the same methods (MAGUS and FastTree):

For running WITCH to separate the backbone and queries, and to produce the backbone alignment and tree,

```
python3 witch.py -t 16 -i $unaln_seqs -d $work_dir
```

For running MAGUS (0.1.0b2):

```
python3 magus.py -np 16 -i $unaln_seqs -d $work_dir -o $output_aln_path
```

For running FastTree (v2.1.11 SSE3):

```
FastTreeMP -lg $bb_aln > $bb_tre # for AA data
FastTreeMP -nt -gtr $bb_aln > $bb_tre # for NT data
```

### 3 Dataset Availability

We list the sources of our datasets here with links to the download pages

- 1000M-HF [6]
- RNASim-500bp [3]
- CRW [1] (5S series (preprocessed by [4]), 16S series)
- 10AA [3]
- HomFam [5]

[4] Chengze Shen, Paul Zaharias, and Tandy Warnow. Magus+ehmms: Improved multiple sequence alignment accuracy for fragmentary sequences, 2021.

[5] Fabian Sievers, Andreas Wilm, David Dineen, Toby J. Gibson, Kevin Karplus, Weizhong Li, Rodrigo Lopez, Hamish McWilliam, Michael Remmert, Johannes Söding, Julie D. Thompson, and Desmond G. Higgins. Fast, scalable generation of high-quality protein multiple sequence alignments using Clustal Omega. *Molecular Systems Biology*, 7:539, October 2011.

[6] Vladimir Smirnov and Tandy Warnow. Phylogeny Estimation Given Sequence Length Heterogeneity. *Systematic Biology*, 70(2):268–282, March 2021.
